# Orphan G protein-coupled receptor, GPR37L1: pharmacological toolbox empty once again

**DOI:** 10.1101/2020.09.11.290486

**Authors:** Tony Ngo, Brendan P. Wilkins, Sean S. So, Peter Keov, Kirti K. Chahal, Angela M. Finch, James L.J. Coleman, Irina Kufareva, Nicola J. Smith

**Affiliations:** Skaggs School of Pharmacy and Pharmaceutical Sciences, UC San Diego, La Jolla, CA, United States; Molecular Pharmacology Laboratory, Victor Chang Cardiac Research Institute, Darlinghurst, NSW, Australia; Orphan Receptor Pharmacology Laboratory, School of Medical Sciences, UNSW Sydney, Kensington, NSW, Australia; G Protein-Coupled Receptor Laboratory, School of Medical Sciences, UNSW Sydney, Kensington, NSW, Australia

## Abstract

Orphan G protein-coupled receptors (GPCRs) are largely intractable therapeutic targets, owing to the lack of chemical tools for exploring their pharmacology. The discovery of such tools, however, is hampered by a number of unknowns, such as effector coupling and appropriate positive controls. In our 2017 Nature Chemical Biology paper^1^, we developed a computational chemical tool discovery approach called GPCR Contact-Informed Neighboring Pocket (GPCR-CoINPocket). This method predicted pharmacological similarity of GPCRs in a ligand- and structure-independent manner, to enable the discovery of off-target activities of known compounds at orphan GPCRs and hence the identification of so-called surrogate ligands. Our orphan GPCR target for prospective surrogate ligand discovery efforts was GPR37L1, a brain-specific receptor linked to cerebellar development^2^ and seizures^3^. We had previously demonstrated that GPR37L1 constitutively coupled to Gαs and generated ligand-independent increases in intracellular cAMP^4§^. Thus, the inverse agonist activities of computationally predicted surrogates were tested in the cAMP response element luciferase (CRE-luc) reporter gene assay in human embryonic kidney (HEK293) cells expressing either vector control or what we thought was untagged GPR37L1 in pcDNA3.1. However, we recently discovered that the GPR37L1 construct used in that study was incorrect: instead of pcDNA3.1, it carried the receptor inserted backwards into a yeast p426GPD vector (hereafter referred to as p426-r37L1). Here, we correct the cloning error and describe our subsequent unsuccessful efforts to re-test the computationally predicted GPR37L1 ligands (triggering an author-initiated retraction of^1^).

**Note:** We, the authors, are working with the Nature Chemical Biology Editors to retract our 2017 paper ‘Orphan receptor ligand discovery by pickpocketing pharmacological neighbors’^1^. The present manuscript is under review at Nature Chemical Biology as a Matters Arising accompaniment to the anticipated author-initiated retraction. We initiated the steps towards the retraction upon discovering a regrettable cloning error that put into question the in vitro findings reported in^1^. This action was unanimously agreed upon by all authors. The computational aspects of the original manuscript^1^ are unaffected by this error.

---

In the CRE-luc assay, p426-r37L1 had considerably elevated transcriptional reporter activity when compared to pcDNA3.1, which had led to the interpretation that GPR37L1 constitutively activated Gαs and promoted cAMP production; however this elevated signal is also evident for empty p426GPD vector (**Fig1a**). In contrast, neither the corrected wild type GPR37L1 construct, nor GPR37L1 C-terminally tagged with enhanced yellow fluorescent protein (GPR37L1-eYFP), were different from baseline. DNA titration of p426-r37L1 in an upstream orthogonal, real-time cAMP assay with the BRET-based CAMYEL biosensor^5^, produced results that were no different from pcDNA3.1 vector alone, while titration with the positive control β_2_-adrenoceptor resulted in a concentration-dependent signal increase (**Fig1b**). The titration with pcDNA3.1-GPR37L1 also produced no increase in CAMYEL signal, in agreement with the CRE-luc assay and consistent with lack of constitutive Gαs-directed activity. Receptor expression by the corrected construct was confirmed by Western blotting (**Fig1c**). Together these results suggest that the aberrant signal generated by p426-r37L1 was due to interference caused by the p426-GPD vector and specific to the CRE-luc assay, and that the real GPR37L1 is not constitutively active towards Gαs.

**Figure 1:**
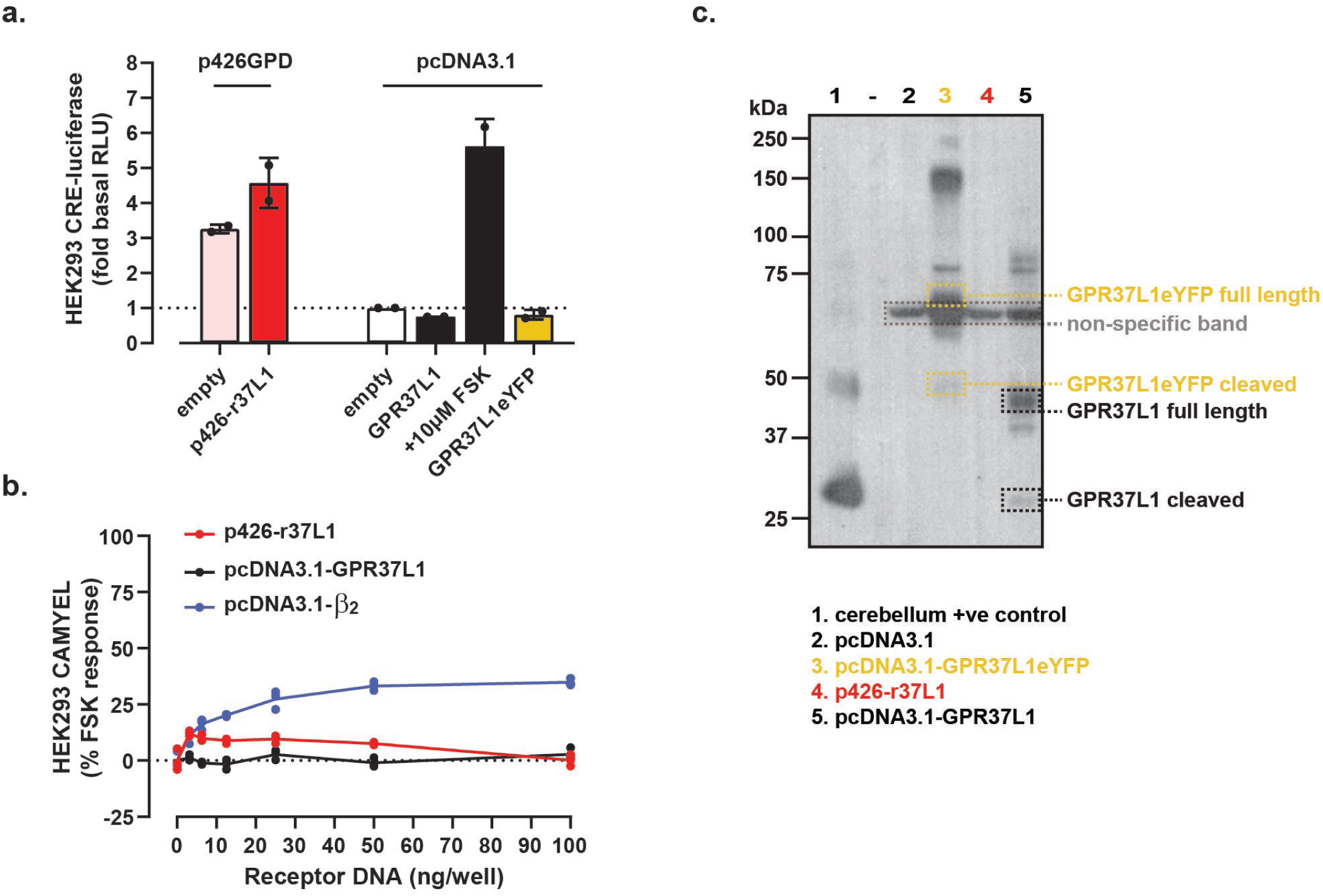
The corrected GPR37L1 construct does not have the apparent Gαs constitutive activity seen with p426-r37L1. **(a-b)** Examination of the effects of p426-r37L1 on reporter signal in **(a)** CRE-luc and **(b)** CAMYEL assays. HEK293 cells were transiently co-transfected with the CRE-luc reporter or the CAMYEL BRET biosensor together with incorrect or corrected GPR37L1 construct or β_2_-adrenoceptor, as indicated. ‘Empty’ refers to empty plasmid DNA. For CRE-luc assay, data represent n=2 biological replicates performed in triplicate. For CAMYEL assay, data represent n=1 with technical replicates shown as circles. RLU, relative light units; FSK, forskolin. **(c)** Western blot for transient HEK293 cellular expression of r37L1, corrected GPR37L1 or controls, as indicated. Cerebellum from a male C57BL/6J mouse was used as a positive control. Dashed lane contained protein ladder only. Image is n=1.

Next, we focused on the computationally predicted compounds that were reported in^1^ to be inverse agonists of Gαs-directed constitutive activity of GPR37L1. Because the corrected construct did not generate such activity (**Fig1a-c**), their inverse agonism could not be confirmed. The compounds were not Gαs-directed agonists of GPR37L1 (**Fig2a-b**) and their antagonism could not be tested for the lack of a control agonist; the previously reported GPR37L1 agonist prosaptide TX14A^6,7^ had no activity towards Gαs (**Fig2a-c**). These findings held true in two cellular contexts, HEK293 (**Fig2a-b**) and CHO cells (**Fig2c**), and in two Gαs-cAMP pathway assays, CRE-luc (**Fig2a**) and CAMYEL (**Fig2b-c**). A positive control, the constitutively active Gαs-coupled β_2_ adrenoceptor, behaved as expected. The “lead” compound from^1^, SHA68, was also tested in the receptor-proximal AP-TGFα sheddase assay^8^ in FlpIN TREx HEK293 cells inducibly expressing GPR37L1-eYFP and co-transfected with Gαq/s (**Fig2d**); this chimeric G protein directs Gαs-coupled receptor signaling through the Gαq/TGFα pathway. Gαs-directed agonism was apparent for the positive control, isoproterenol, in β_1_ adrenoceptor-expressing cells, but not for SHA68 at GPR37L1 (**Fig2d**). Altogether, this suggests that the compounds reported in^1^ do not activate Gαs-mediated signaling by GPR37L1, whereas their ability to inhibit such signaling could not be tested.

**Figure 2:**
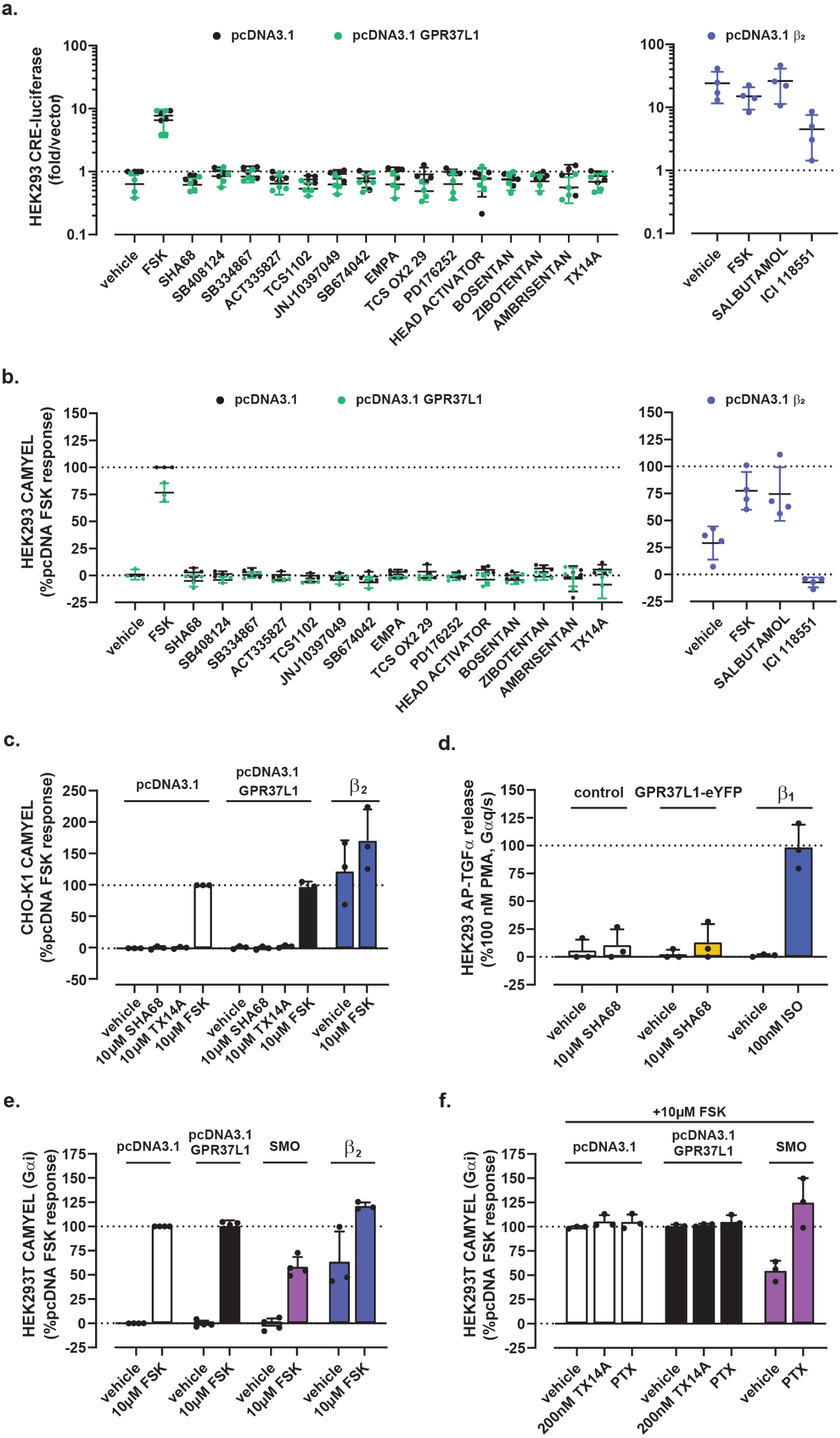
The corrected GPR37L1 construct does not display constitutive or ligand-induced activity in any assays examined. Examination of constitutive and ligand-induced cAMP elevation in HEK293 cells transfected with or without the corrected pcDNA3.1-GPR37L1, using **(a)** CRE-luc or **(b)** CAMYEL assays. β_2_-adrenoceptor +/- 10 μM salbutamol was used as a positive control for constitutive cAMP signaling. Data represent n=3-4 assays performed in triplicate and final ligand concentrations were 10 μM. **(c)** CAMYEL assay performed as in (b) using CHO-K1 cells to control for cellular background; n=3 performed in triplicate. **(d)** Ligand-induced Gαs coupling was probed using the AP-TGFα-sheddase assay in stable FlpIN TREx HEK293 cells inducibly expressing GPR37L1-eYFP, transiently transfected with AP-TGFα and chimeric Gαq/s. β_1_ adrenoceptor stimulated with isoproterenol (ISO) was used as a positive control. Data represent n=3 independent experiments, each performed in triplicate. **(e-f)** Constitutive suppression of FSK-stimulated cAMP was monitored using the CAMYEL BRET biosensor in HEK293T cells transiently co-transfected with CAMYEL and either the corrected GPR37L1 plasmid, Smoothened, or β_2_ adrenoceptor. Gαi3 was transiently co-transfected with GPR37L1 and Smoothened. Data represent n=3-4 assays performed in triplicate. All data panels are presented as mean +/- standard deviation.

Next, because the inchoate GPR37L1 literature contains conflicting reports of Gαs vs Gαi coupling specificity of this receptor^3,4,6,7,9,10^, we turned our attention to the Gαi pathway. Unlike the positive control Smoothened receptor^11^, GPR37L1 showed no Gαi-directed constitutive activity as indicated by the lack of constitutive, pertussis-toxin-sensitive suppression of forskolin-stimulated cAMP in the CAMYEL assay in HEK293T cells (**Fig2e-f**). No cAMP suppression was observed following cell pretreatment with prosaptide TX14A either (**Fig2f**). This lack of constitutive GPR37L1 activity or activation by the previously reported agonist precluded investigations of inverse agonism or antagonism of the compounds from^1^ in the Gαi pathway. Altogether, our data suggests that the pharmacological toolbox for GPR37L1 is once again empty.

So where does this leave the computational GPCR-CoINPocket methodology developed in^1^? Despite the mishap with GPR37L1, the methodology remains a powerful tool for understanding pharmacological similarity of GPCRs solely from their sequences. In^1^, the method was retrospectively benchmarked using the ChEMBL database and curated literature examples. Prospectively, GPCR-CoINPocket correctly predicted the pharmacological similarity of the orphan receptor GPR31 to the hydroxycarboxylic acid (HCA) receptor family; the prediction was later independently confirmed by the experimental identification of ligands that GPR31 shared with the HCA_1_ receptor^12^. The CoINPocket methodology was also used to identify new ligand pairings in the evolutionarily unrelated protein lysine methyltransferase protein family^13^. Beyond the search for pharmacological similarities, the GPCR-CoINPocket approach helped improve template selection for accurate GPCR homology modeling^14^ and delineate the common activation mechanism in Class A GPCRs^15^. These examples illustrate the breadth of impact of GPCR-CoINPocket at the intersection of computational and experimental biology.

What of GPR37L1? Orphan GPCR research is frustrated by the absence of pharmacological tools. Indeed, a common shortcoming in all functional GPR37L1 studies, including our own, is that the limited armamentarium forces researchers to infer specificity from indirect and sometimes inconclusive data. For example, phenotypic or second messenger amplification assays with the read-out many steps removed from receptor activation are vulnerable to non-specific signal interference, as was the case for p426-r37L1^1,4^, or to interference from the endogenous receptor context^6,7^. These are common challenges for orphan research^16,17^, which nevertheless should not deter us from continuing to define the pharmacology and physiology of GPR37L1.

## Acknowledgements

We would like to thank Dr Robert M. Graham from the Victor Chang Cardiac Research Institute, Australia, for helpful discussions, Mr Theodore Nettleton, UNSW Sydney, for assistance with identifying the cloning error, and Dr Asuka Inoue, Tohoku University, for providing the AP-TGFα constructs and helpful advice. Funding - TN (NHMRC CJ Martin Early Career Fellowship 1145746), IK (NIH R01 AI118985 and R01 GM117424), NJS (National Heart Foundation Future Leader Fellowship 101153).

## Author Contributions

NJS, JLJC, SSS and BPW discovered the cloning error. NJS, IK, TN, BPW, SSS and JLJC devised the experimental plan for correcting the scientific record; TN, BPW, SSS, PK and KKC performed experiments; AMF assisted with experimental design; NJS, IK and TN wrote the manuscript with input from all of the authors.

## Competing Interests

The authors declare no competing interests.

Correspondence and requests for materials should be addressed to

## Methods

### Compounds and reagents

Tissue culture and Western blot reagents were purchased from Sigma-Aldrich (St. Louis, MO) or Life Technologies (Carlsbad, CA) unless otherwise indicated. Compound suppliers are listed in Ngo et al.^1^.

### Plasmids

Full length untagged human GPR37L1 with the native signal peptide, in pcDNA3.1, was synthesized by Genscript. p426-r37L1 (incorrectly presumed to be pcDNA3.1-GPR37L1 in^1^) contained full-length WT GPR37L1 inserted backwards into a yeast p426GPD vector. pcDNA3.1-GPR37L1-eYFP was described in^4^ and human β_2_ adrenoceptor in pcDNA3.1 was from cDNA Resource Centre (cat #AROB200000). For BRET-based cAMP reporter assays, pcDNA3.1-CAMYEL was used^5^. pGEN-mSmo was a gift from Philip Beachy (Addgene plasmid # 37673,^19^), and modified with a 2xStop codon following the receptor C-terminus. Rat Gαi3 in pcDNA3.1+ was a gift from Pradipta Ghosh (UC San Diego). All plasmids used in TGFα shedding assays^8^, except GPR37L1, were a generous gift from Asuka Inoue (Tohoku University).

### CRE-luciferase cAMP reporter assay

CRE-luciferase reporter gene assays were performed as previously described^20^ with minor modifications. Briefly, HEK293 cells were seeded onto transparent 96-well plates at a density of 20,000 cells/well in DMEM containing 10% FBS and 1% penicillin/streptomycin solution. The next day, CRE-reporter DNA (0.04 μg/well) (Promega) was co-transfected with either pcDNA3.1, corrected pcDNA3.1-GPR37L1, or the erroneous p426-r37L1 construct (0.04 μg/well), or pcDNA3.1-β_2_ adrenoreceptor (0.02 μg/well; positive control) using Lipofectamine LTX. The following day, compounds diluted in DMEM were added to the cells (10 μM final concentration). After approximately 21 hr, cells were washed once with PBS before lysis with 40 μl Passive Lysis Buffer (Promega). For ligand screening experiments, lysates were frozen and thawed the next day to allow uniform processing of samples. To measure luciferase activity, 4 μL of lysate was transferred to a white OptiPlate-384 in duplicate. Following injection of 10 μl Luciferase Assay Reagent II (Promega), luminescence was detected immediately using a PHERAstar FSX plate reader (BMG-Labtech).

### Real-time cAMP measurements with the CAMYEL Bioluminescence Resonance Energy Transfer (BRET) biosensor

For Fig1b and Fig2b, HEK293 cells were seeded onto white 96-well plates at 20,000 cells/well. The next day cells were transfected using Lipofectamine LTX according to manufacturer’s instructions. For the DNA titration experiment (Fig1b), CAMYEL (0.02 μg/well) and construct of interest (0-0.1 μg/well) were co-transfected. For the experiments in Fig2b, 0.02 μg CAMYEL and 0.02 μg of receptor DNA were co-transfected per well. The following morning, cells were washed with 200 μL/well Hank’s Balanced Salt Solution (HBSS) and then incubated in 85 μl HBSS at 37°C for 30 min. Luminescence was initiated by addition of a mixture of 2 μM coelenterazine-h (Promega) and 40 μM 3-isobutyl-1-methylxanthine (IBMX; final concentrations listed) in HBSS; kinetic readings were taken for 5 min baseline and then for 15 min after addition of ligands, all at 37°C, using the BRET1 filter set on a PHERAstar plate reader (BMG-Labtech). Data presented are the 15 min endpoint read. Experiments in Fig2c were performed similarly except CHO-K1 cells were maintained in Ham’s F12 media containing 10% FBS and 1% penicillin/streptomycin solution. For Fig2e-f, HEK293T cells were plated at 700,000 cells/well in a 6-well plate and transfected next day with 1.0 μg/well CAMYEL, 0.5 μg/well Gαi3 (for GPR37L1 and SMO only), and 1.0 μg/well of the indicated receptors using TransIT-X2 transfection reagent (Mirus Bio); the total transfected DNA was normalized to 3.0 μg/well for all wells using empty pcDNA3.1. The following day, cells were lifted by gentle pipetting and re-plated in a 96-well plate at 50,000 cells/well. On day 4, cells were serum-starved in assay buffer (1X HBSS, 20 mM HEPES, 0.05% BSA) for 1 hr and then IBMX and coelenterazine-h were added to each well to the final concentrations of 100 μM and 10 μM, respectively, for 10 min. Emission intensity was recorded for 5 min using 460 nm and 530 nm filters on the EnVision or Victor X Light plate reader (PerkinElmer). Next, 10 μM FSK was added to each well and emission intensities at 460 nm and 530 nm were continuously read for another 20 min. In Fig2f, 200 nM TX14A was added after 5 min, and continuously read for 10 mins before FSK addition. PTX (200 ng/μL) was added at least 3 h prior to serum starvation in assay buffer. Inverse BRET (1/BRET) was calculated as the window-averaged ratio of emission intensity at 460 nm and 530 nm at 5 min post FSK addition, normalized to FSK response from the pcDNA3.1-transfected cells in the same experiment.

### AP-TGFα sheddase assay

AP-TGFα sheddase assays were performed as described previously^8^. Briefly, GPR37L1-eYFP expression was induced in stable FlpIN TREx HEK293 cells using 100 ng/μL doxycycline. Transient transfections of β_1_ adrenoceptor, AP-TGFα and the Gαq/s chimera were performed using Lipofectamine LTX. Cells were stimulated with 10 μL/well of either isoproterenol for β_1_-adrenoceptor conditions or SHA68 for GPR37L1- eYFP conditions to achieve final concentrations of 100 nM or 10 μM, respectively. Phorbol-12-myristate-13-acetate (PMA, 100 nM) served as a positive control. Absorbance at 405 nm was measured using a PHERAstar FSX plate reader (BMG-Labtech).

### Western blot

HEK293 cells were seeded onto 6-well plates at a density of 400,000 cells/well. The next day, cells were transfected with pcDNA3.1, pcDNA3.1-GPR37L1-eYFP, p426-r37L1, or pcDNA3.1-GPR37L1 (0.4 μg/well) using Lipofectamine LTX. After 24 hr, cells were washed once with ice-cold PBS and lysed with ice-cold RIPA buffer containing cOmplete™ Protease Inhibitor Cocktail (Roche Diagnostics). For murine cerebellar samples, frozen tissue was homogenized in RIPA buffer containing cOmplete™ Protease Inhibitor Cocktail on ice using a POLYTRON homogenizer. The cell and tissue lysates were centrifuged at 18,000 g for 10 min and the resulting supernatant retained. Total protein concentrations were determined using a BCA assay. Loading samples were prepared by mixing cell lysates with NuPAGE LDS sample buffer containing 100 mM dithiothreitol, and heated for 15 min at 65°C. 10 μg of each sample was loaded on a NuPAGE 4-12% bis-tris polyacrylamide gel and separated over 1 hr at 187V. Proteins were transferred onto a polyvinylidene fluoride membrane (Merck Millipore), followed by blocking in Tris-buffered saline containing 0.1% Tween-20 (TBST) and 5% skim milk for 1 hr. Membranes were probed with anti-GPR37L1 primary antibody (1:1000, sc-164532, Santa Cruz Biotechnology) at 4°C overnight, followed by anti-goat horseradish peroxidase-conjugated secondary antibody (1:10,000, Invitrogen) for 1 hr. Chemiluminescence was detected with Clarity™ (Bio-Rad, USA) chemiluminescence substrate using X-ray film (Fujifilm).

## Footnotes

§ note that this paper has already been retracted due to the use of the affected construct, again in the CRE-luc assay. However, this paper also demonstrated GPR37L1 coupling to Gαs in yeast and cerebellar slice cultures by ELISA: these experiments are unaffected by the error and their results remain valid^18^.

